## Supplementary Material for "Guanidine Production by Plant Homoarginine-6-hydroxylases"

To the manuscript

**The supplementary material comprises:**

- Figures S1 to S13
- Tables S1 to S3
- SI References

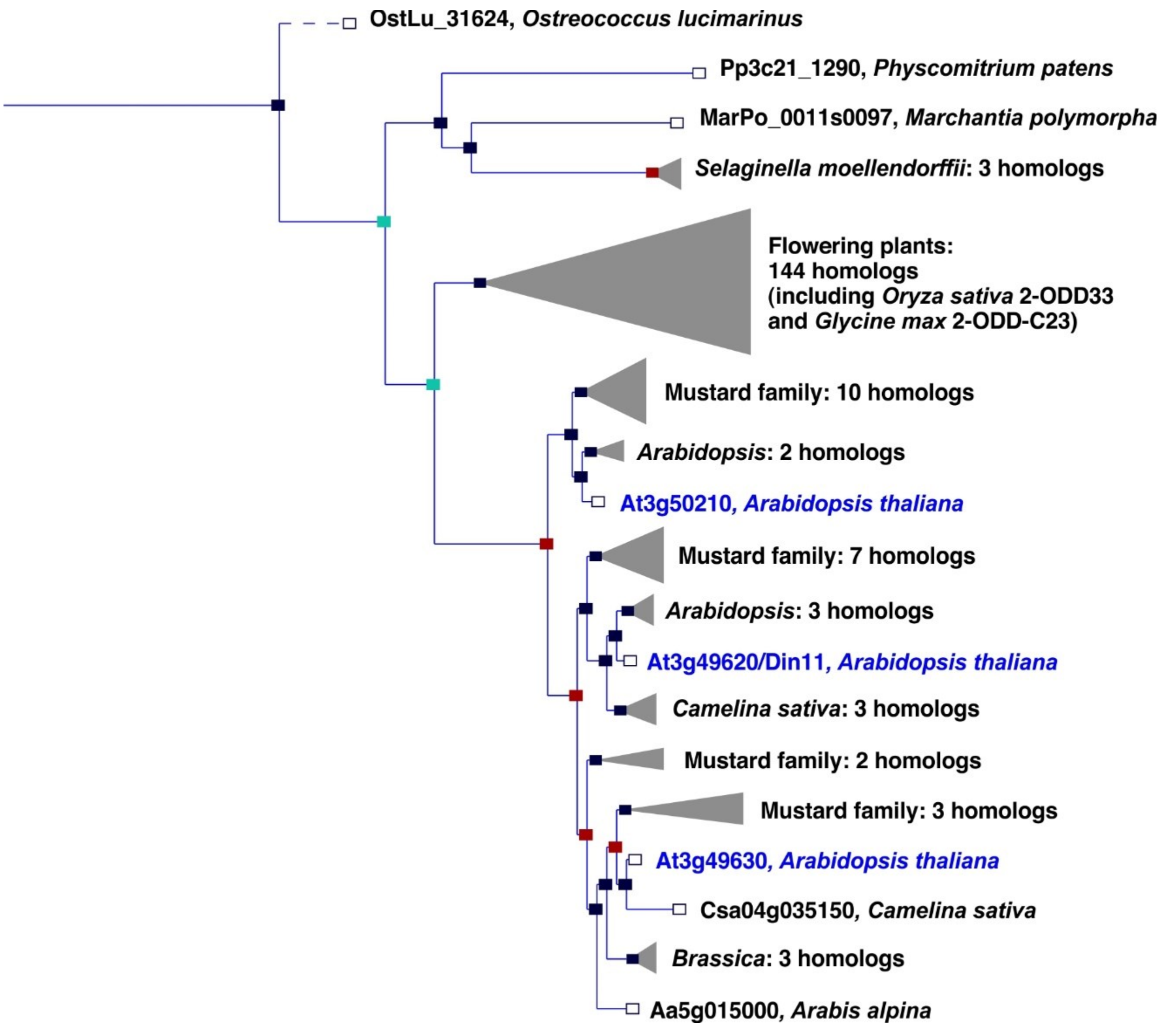


**Fig. S1:** Phylogeny of the 2-ODD clade C23. Excerpt of the gene tree for *Din11* from EnsemblPlants (https://plants.ensembl.org, accessed Feb. 2023). The length of the horizontal blue lines is proportional to the number of amino acid exchanges except the dashed line, which is shortened by a factor of 10. Blue squares indicate speciation nodes, red squares indicate duplication nodes, green squares indicate ambiguous nodes, and open squares indicate species nodes. Grey triangles indicate collapsed branches. The three 2-ODD-C23 paralogues from *Arabidopsis thaliana* are highlighted by blue lettering.

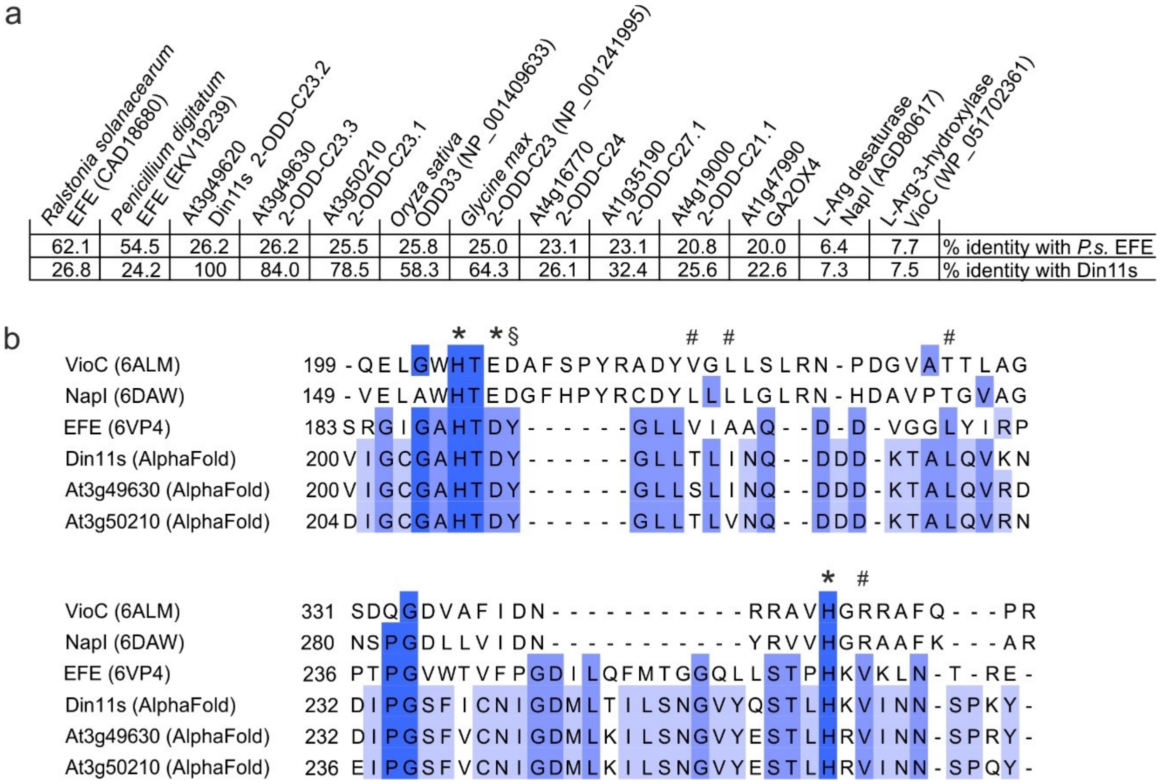


**Fig. S2:** Sequence comparison of structurally or functionally related 2-ODD proteins. (a) Sequence identity between *Pseudomonas savastanoi* EFE (*P.s.* EFE), Din11s and selected other members of the 2-ODD superfamily derived from a multiple sequence alignment containing further plant and bacterial sequences. BLAST searches with NapI or VioC do not produce significant hits in Arabidopsis. Gibberellin 2-oxidase 4 (GA2OX4) has the highest degree of similarity with *P.s.* EFE among the functionally characterised Arabidopsis 2-ODDs. NCBI protein accession numbers are given in brackets. (b) Excerpt of a sequence alignment derived from a structural superposition of *P.s.* EFE with the structures other 2-ODDs acting on arginine or homoarginine as substrate. The structural alignment was generated in Chimera1.15 and was based only on the residues surrounding the active site, because an overall alignment was not possible. Residues contributing to $Fe^{2+}$-coordination in the active site are marked with *. Residues contributing to 2-oxoglutarate binding in *P.s.* EFE are marked with #. A tyrosine residue that is structurally and functionally important for arginine binding in *P.s.* EFE is marked with §. This tyrosine residue is present in 2-ODD-C23 enzymes but not in any other Arabidopsis 2-ODD sequences.

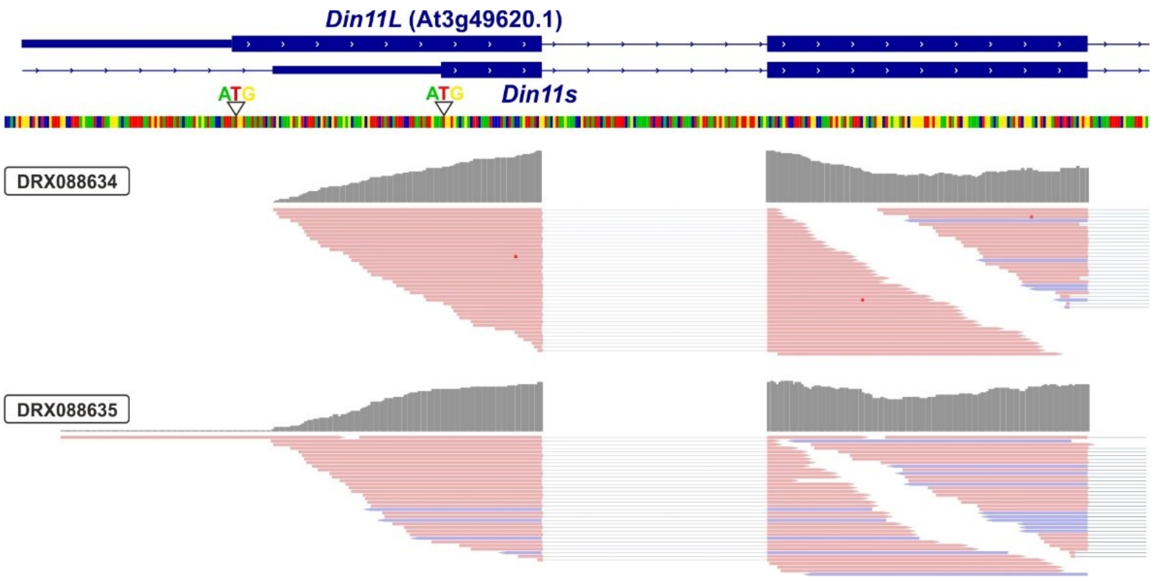


**Fig. S3:** 3′-UTR of *Din11*. Alignment of RNAseq reads (pink or purple lines) from two independent experiments with the genomic sequence (four-coloured line) of Arabidopsis around the transcription start site of *Din11*. The 5′-end of the annotated *Din11* transcript At3g49620.1/*Din11L* is hardly covered by RNAseq reads, which are much more frequent several basepairs downstream of the annotated start codon. The graphical representation of the RNAseq reads was modified from the Arabidopsis RNAseq database (Zhang et al., 2020).

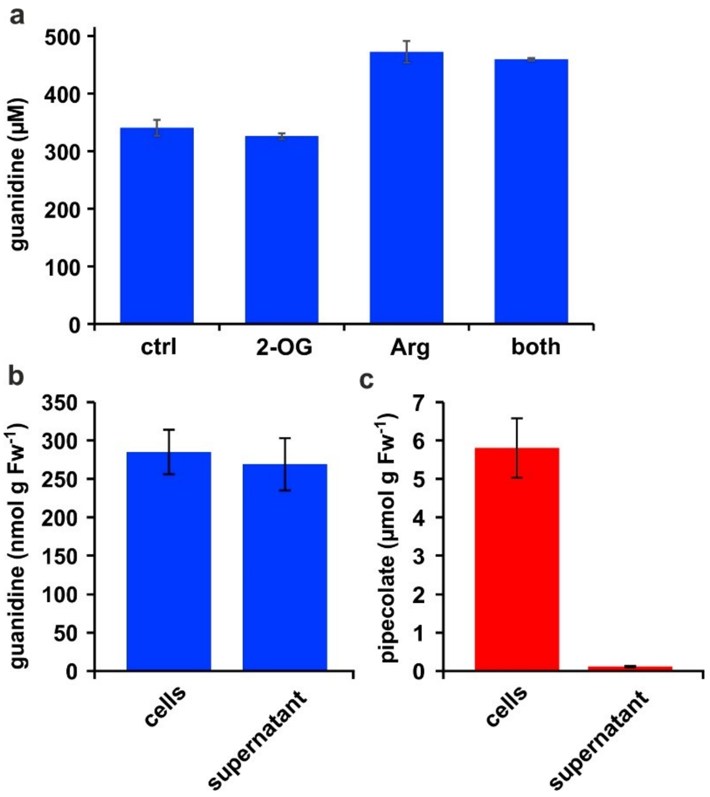


**Fig. S4:** Distribution of metabolites in *E. coli* cultures. (a) *E. coli* cells expressing Din11s were cultivated over night at 18 °C in medium supplemented with either 5 mM 2-oxoglutarate (2-OG), 5 mM arginine (Arg), or a combination of both. The cultures were diluted with a 20-fold excess of methanol containing 10 µM $^{13}C^{15}N$-labelled guanidine as analytical standard. (b), (c) *E. coli* cells expressing soybean 2-ODD-C23 were grown over night at 18 °C in the presence of 1 mM homoarginine. The cells were harvested by centrifugation and lysed in 20 µl (mg Fw)$^{-1}$ methanol containing 10 µM labelled guanidine. The supernatant was also mixed with a 20-fold excess of methanol with 10 µM labelled guanidine. Guanidine and pipecolate were quantified by LC-MS. The columns are the average ± SD of samples from three independent cultures. The experiments were repeated with consistent results.

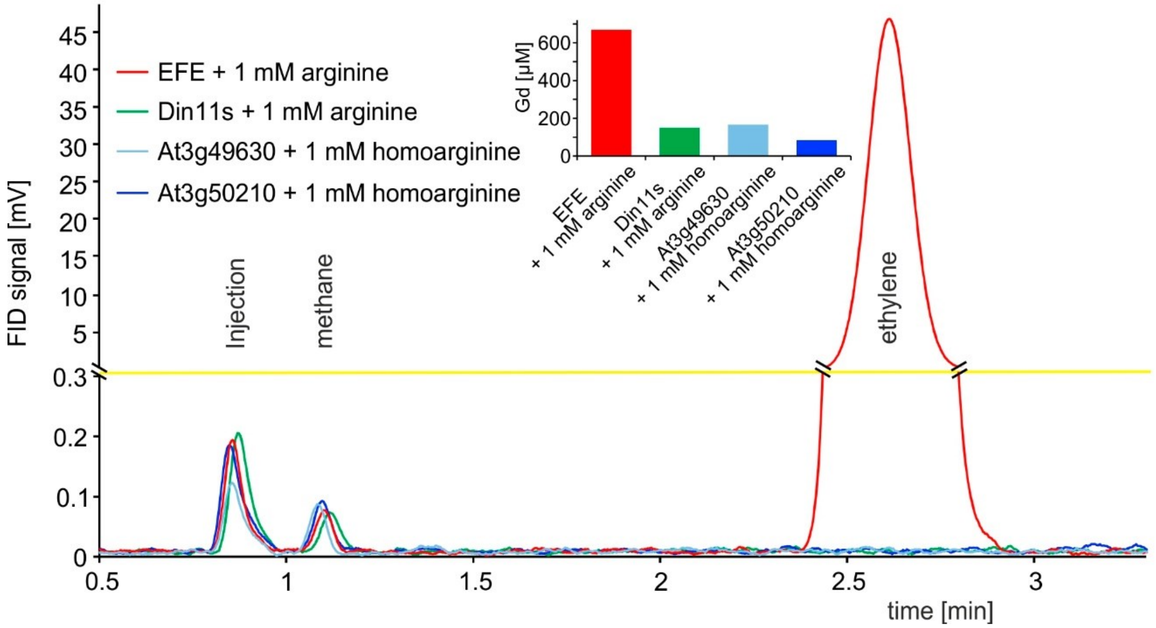


**Fig. S5: 2-ODD C23 isoforms do not produce ethylene.** Flame ionization detector (FID) traces of samples from the headspace of *E. coli* cultures overexpressing *Pseudomonas savastanoi* EFE, Din11s, At3g49630, or At3g50210 and supplemented with 1 mM arginine or homoarginine as indicated in the figure. One ml of culture headspace was injected into a gas chromatograph (SGI 8610C, SRI Instruments, Los Angeles, CA, United States) equipped with a 3 m HayeSep-D column (80 °C, carrier gas $N_2$). Methane from the ambient air and ethylene produced by the bacteria were detected with a flame ionization detector. The software PeakSimple v4.44 was used to record the chromatograms. The detector output was smoothed by averaging 9 measurements recorded at 3 Hz. Inset: Quantification of guanidine in the cultures by LC-MS was used to confirm that the enzymes had been active. All data are from single measurements and consistent results were obtained in independent replicates.

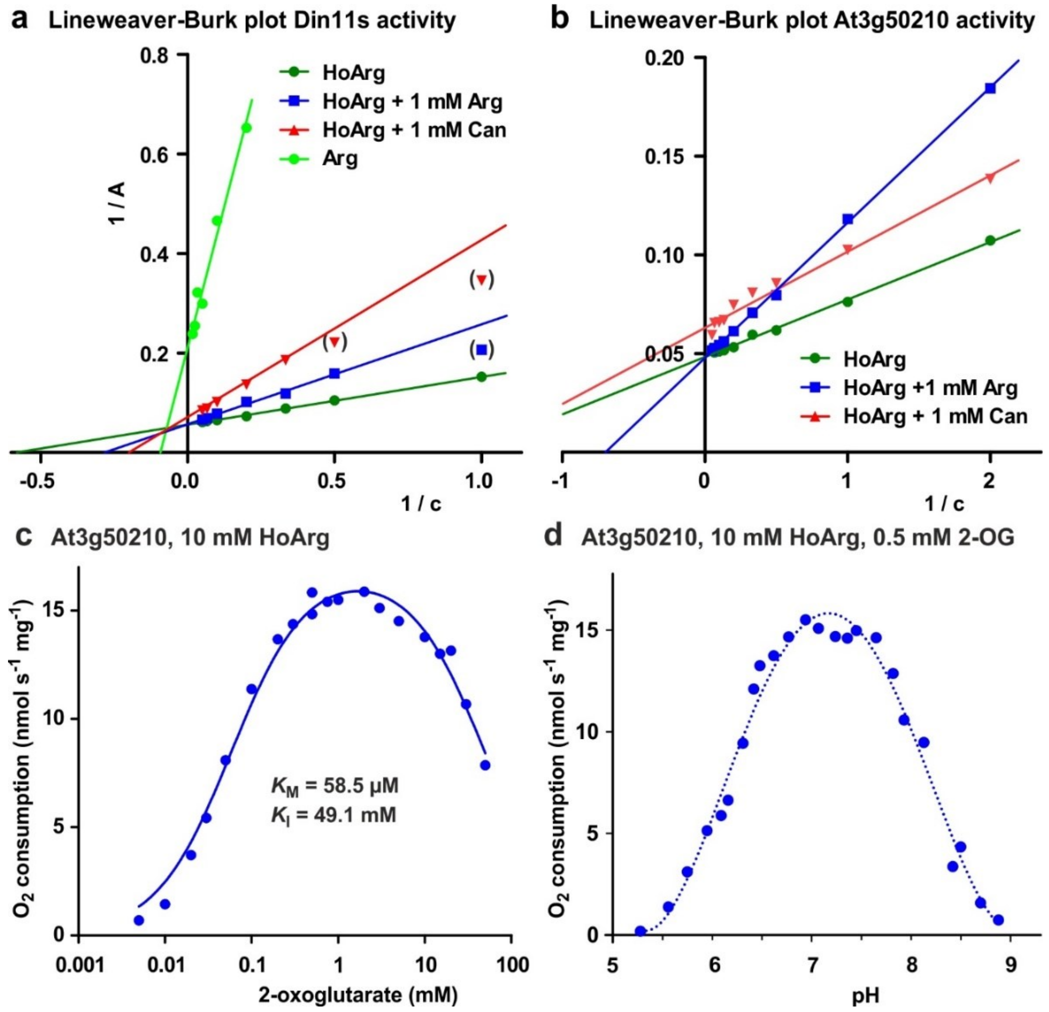


**Fig. S6:** Biochemical characterization of 2-ODD-C23 isoforms. (a), Lineweaver-Burk plot of the activity data for Din11s from Fig. 2a. Values in brackets are very low activities at low concentrations of homoarginine (HoArg) in presence of 1 mM arginine (Arg) or canavanine (Can) that were excluded from the linear regression. (b), Lineweaver-Burk plot of the activity data for At3g50210 from Fig. 2c. Regression lines intersecting on or very near the Y-axis indicate competitive inhibition. (c) Specific activity of At3g50210 in dependence on the concentration of the co-substrate 2-oxoglutarate was determined at 30 °C. Homoarginine concentration was fixed at 10 mM. The blue line represents the least square fit to the Michaelis-Menten equation with an additional term for substrate inhibition. (d), pH-dependence of the specific activity of At3g50210 in a mixed buffer system with MES, HEPES and CHES. Homoarginine and 2-oxoglutarate (2-OG) concentrations were fixed at 10 mM and 0.5 mM, respectively. The dashed blue line is a polynomic fit with a 4$^{th}$-order polynomial. Dots represent single measurements and independent enzyme preparations gave consistent results.

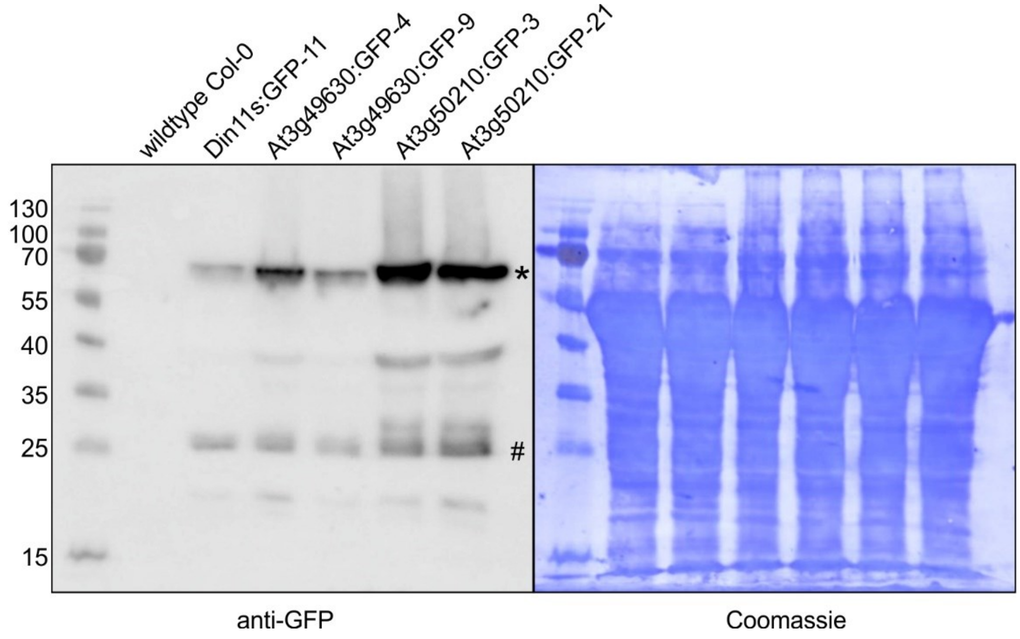


**Fig. S7:** Detection of GFP fusion proteins by Western blot. In soluble protein extracts of rosettes from all transgenic lines expressing 2-ODD-C23:GFP fusion proteins, an anti-GFP antibody detected proteins of the expected size (calculated MW: 68 kDa; *). Minor signals at 27 kDa (#) may represent free GFP generated by partial proteolysis. Staining of the membrane with coomassie brilliant blue was used as loading control.

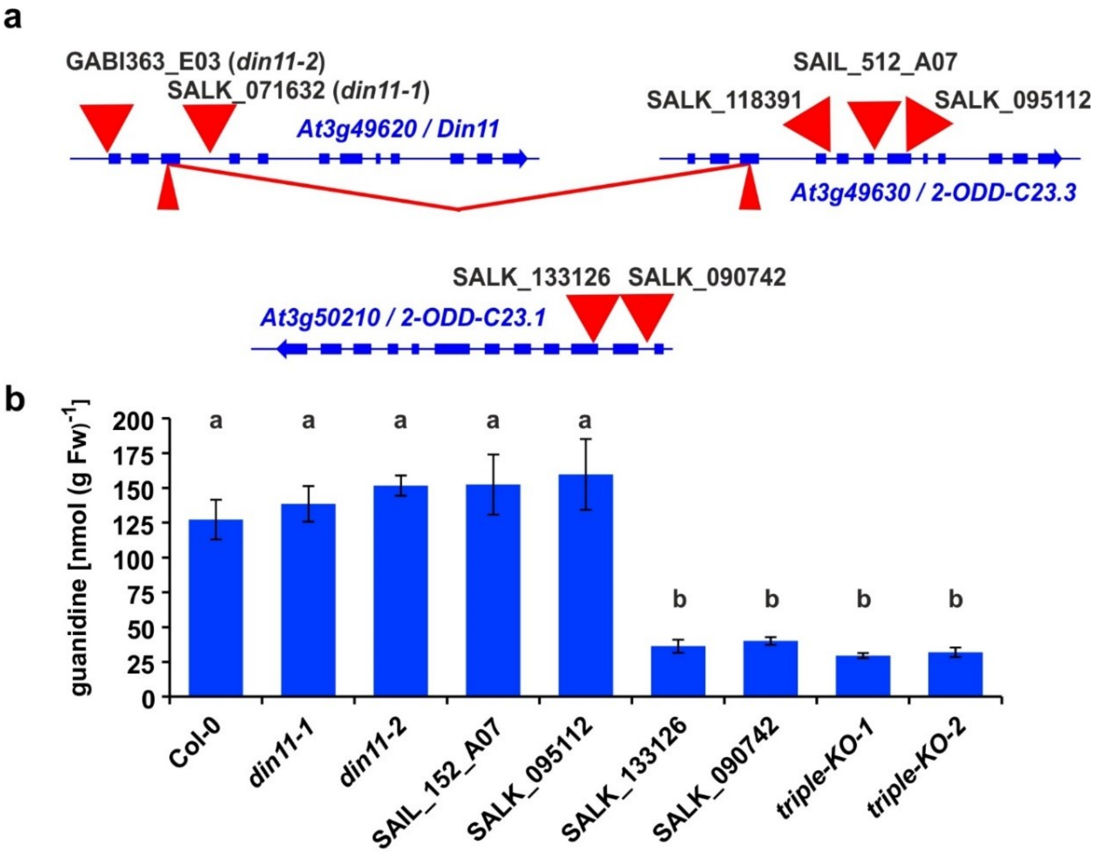


**Fig. S8:** 2-ODD-C23 mutants in Arabidopsis. (a) Schematic representation of the three genes for 2-ODD-C23 isoforms in Arabidopsis. Thick blue lines represent exons, wide red triangles represent T-DNA insertions, narrow red triangles represent CRISPR/Cas9 target sites. The red lines indicate a fusion event between Din11 and At3g49630, in which the DNA fragment between the two CRISPR/CAs9 target sites was lost. (b) Guanidine content in rosettes of single and triple mutant seedlings cultivated under axenic conditions. The seedlings were cultivated for 9 days under long day conditions on half-strength MS medium supplemented with 2% sucrose. Subsequently, they were transferred for one week to plates supplemented additionally with 0.2 mM homoarginine. Columns represent the average ±SD of N = 4 independent samples. Different letters above the columns indicate significant differences ($p < 0.05$) by one-way ANOVA with Bonnferroni correction for multiple testing.

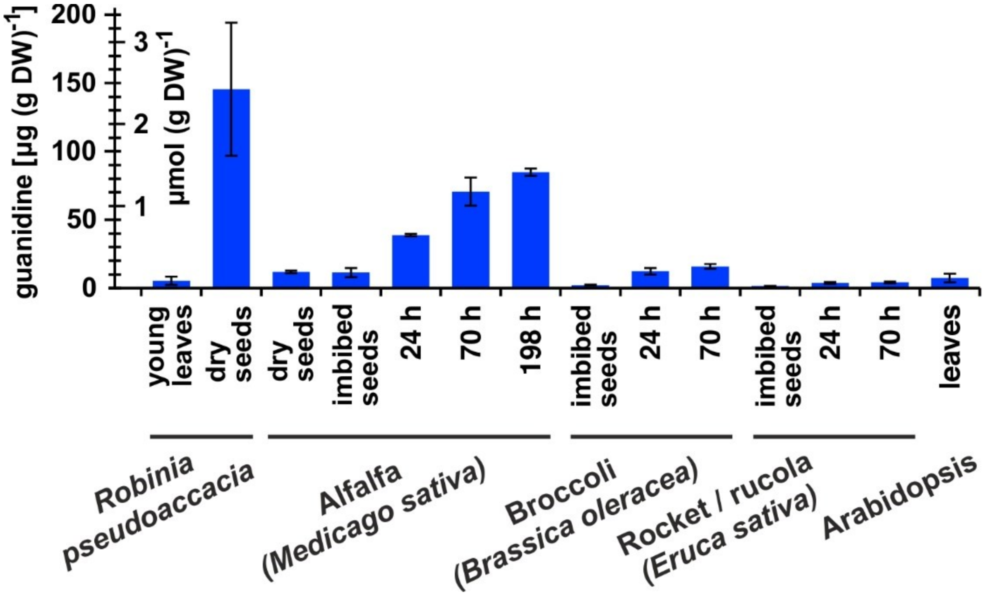


**Fig. S9:** Guanidine content in various plants. Black locust (*Robinia pseudoaccacia*) and Arabidopsis samples were from soil-grown plants. For the other species, seeds were surface-sterilised and germinated on filter paper soaked with dd$H_2O$ for the indicated times. Guanidine was extracted in 80% (v/v) methanol with 10 µM $^{13}C^{15}N$-labelled guanidine as analytical standard and quantified by LC-MS. Columns represent the average ±SD of N = 3 independent samples.

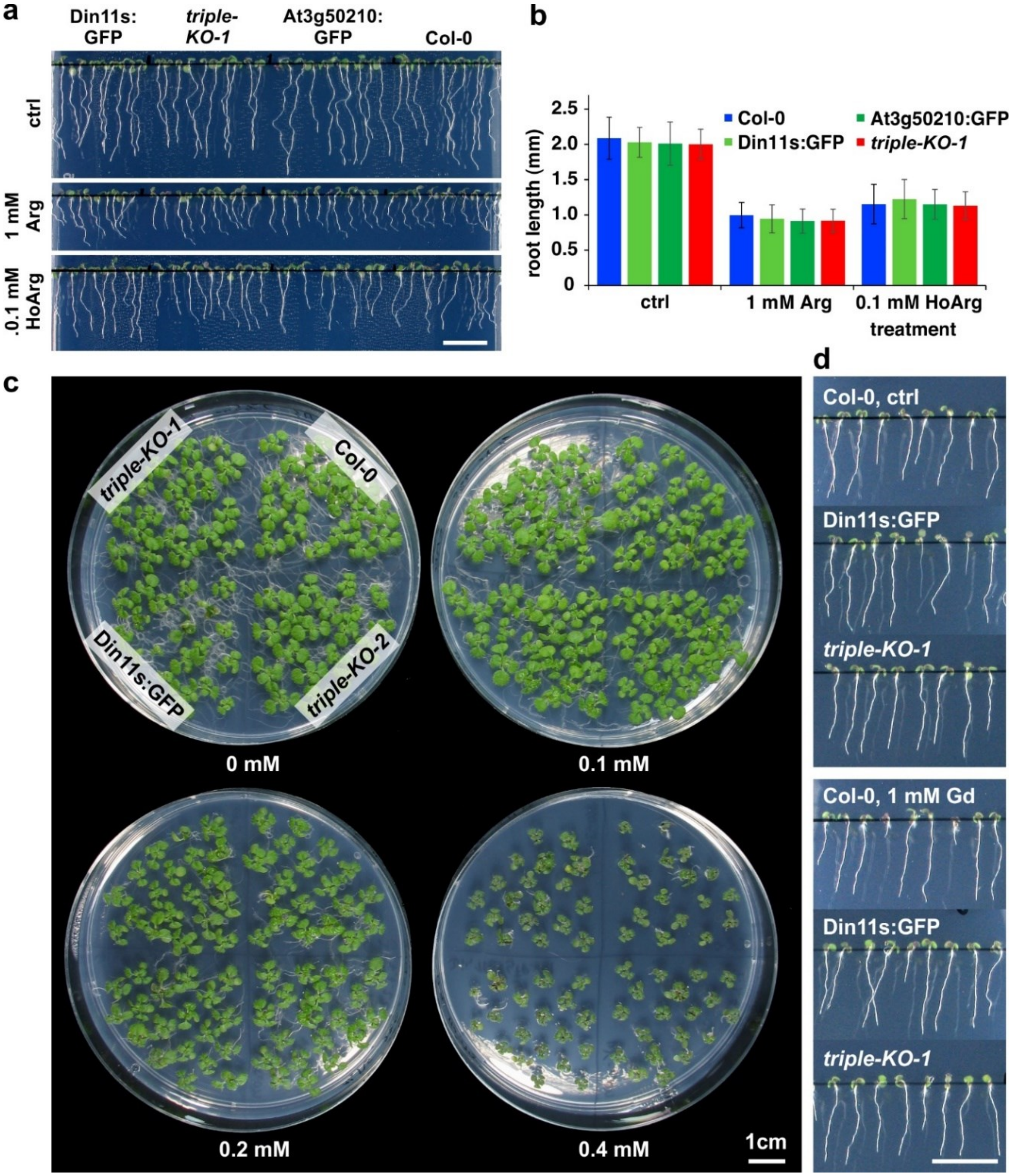


**Fig. S10:** Growth inhibition by homoarginine. (a), (b) Seedlings of WT Arabidopsis plants (Col-0) and 2-ODD-C23 overexpressors or triple mutants were grown for 6 days under long-day conditions on vertical plates containing half-strength MS medium, supplemented with 2% (w/v) sucrose and either arginine (Arg) or homoarginine (HoArg). Columns in (b) represent the average ±SD of N = 40 seedlings. (c) Two-week old seedlings grown on half-strength MS medium with 2% (w/v) sucrose and the indicated concentration of homoarginine. The positioning of the genotypes is identical on each plate. (d) Seven-day-old seedlings grown on vertical plates without or with 1 mM guanidine (Gd). Scale bars in (a), (c) and (d) are 1 cm.

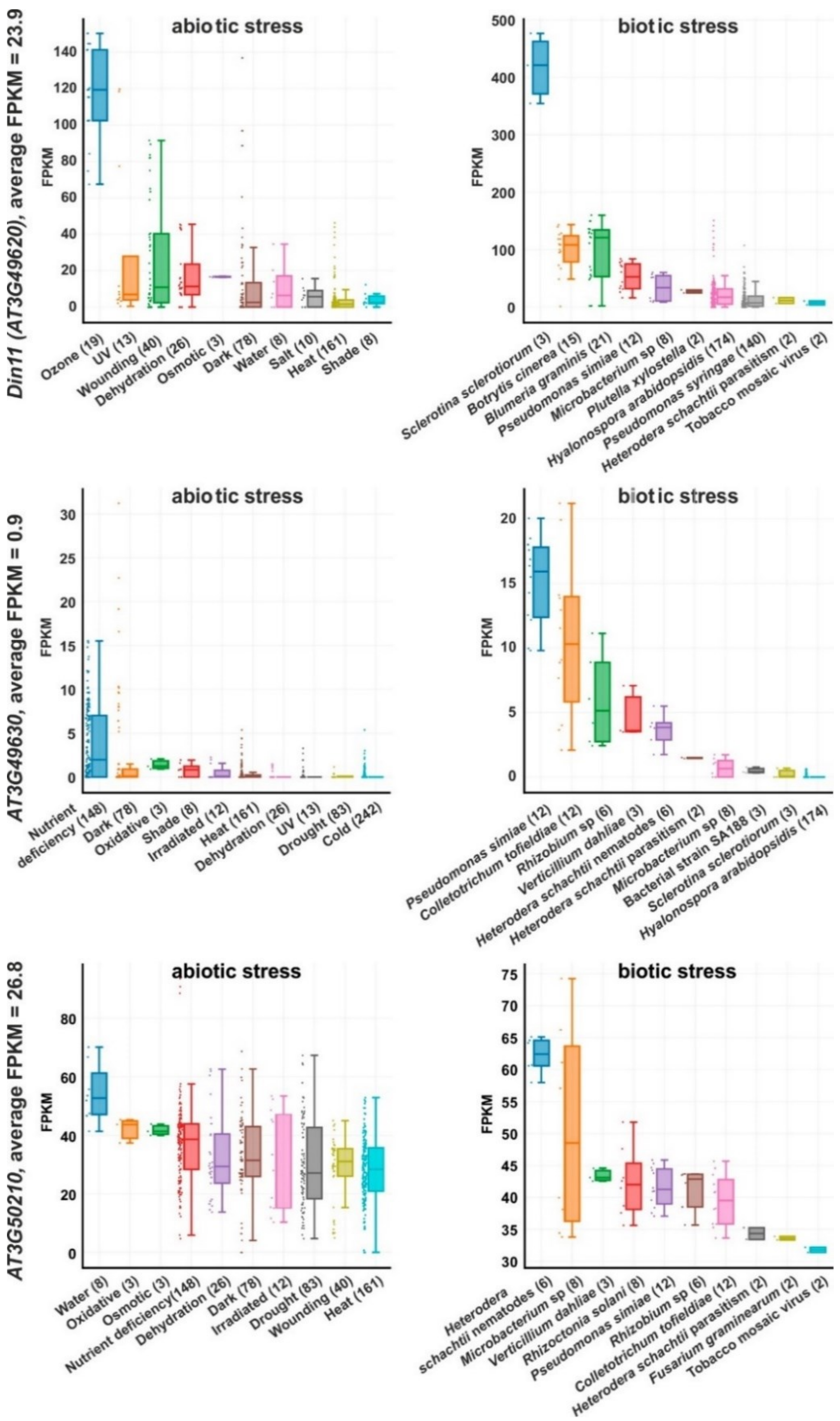


**Fig. S11:** Expression levels of *2-ODD-C23s* under stress. RNAseq coverage of the three Arabidopsis *2-ODD-C23* isoforms under various abiotic and biotic stress conditions. The data plots were modified from the Arabidopsis RNAseq database (Zhang et al., 2020). Numbers in brackets are the number of samples per treatment. FPKM, Fragments per kilobase of transcript per million mapped reads.

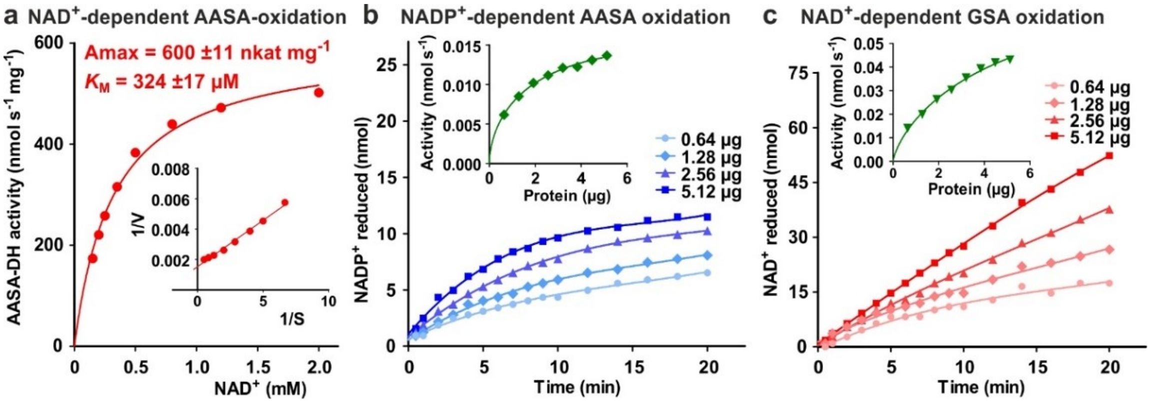


**Fig. S12:** Characterization of ALDH7B4 (At1g54100). (a) Activity of purified recombinant ALDH7B4 with different concentrations of $NAD^+$ at 35 °C. AASA/P6C concentration was fixed at 1 mM. $K_M$ and $A_{max}$ values were obtained by nonlinear regression to the Michaelis-Menten equation (red line). The inset shows the Lineweaver-Burk plot of the same data. (b) AASA/P6C-dependent reduction of $NADP^+$ by different amounts of ALDH7B4. Concentrations of both $NADP^+$ and AASA/P6C were 1 mM. (c) GSA/P5C-dependent reduction of $NAD^+$ by different amounts of ALDH7B4. $NAD^+$ concentration was 1 mM and DL-GSA/P5C concentration was 2 mM. In b and c, the insets show the initial velocity as a function of the amount of protein in the assay. Data points are the average ±SD of technical triplicates. All experiments were repeated with an independent enzyme preparation.

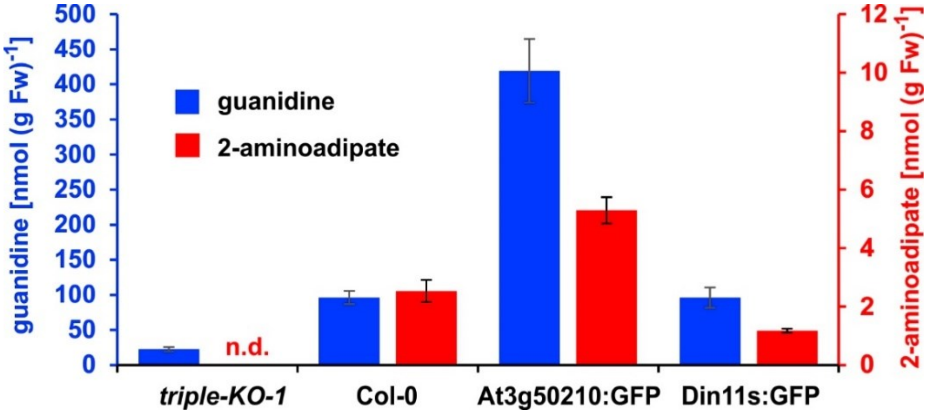


**Fig. S13:** 2-Aminoadipate content after homoarginine feeding. Arabidopsis seedlings were grown for ten days under long-day conditions on half-strength MS medium supplemented with 2% (w/v) sucrose and 5 mM MES-KOH, pH 5.8 before they were transferred for four days to plates additionally containing 0.2 mM homoarginine. Guanidine and 2-aminoadipate were extracted in 80% (v/v) methanol containing 10 µM $^{13}C^{15}N$-labelled guanidine as analytical standard and quantified by LC-MS. Columns represent the average ±SD of N = 4 independent samples. n.d., not detected. The experiment was repeated with consistent results.

**Table S1: Kinetic constants of Arabidopsis 2-ODD-C23 enzymes**

| | Homoarginine variable, 0.5 mM 2-oxoglutarate | | |
|---|---|---|---|
| | Inhibitor (1 mM) | $A_{max}$ (nmol $s^{-1}$ $mg^{-1}$) | $K_M$ (mM) |
| Din11s | none | 13.4 ±3.2 | 1.9 ±0.3 |
| | Arginine | 12.9 ±3.5 | 3.6 ±0.5 |
| | Canavanine | 11.2 ±2.7 | 5.5 ±1.0 |
| At3g49630 | none | 5.6 ±1.2 | 4.6 ±0.2 |
| | Arginine | 5.4 ±1.3 | 4.1 ±0.8 |
| | Canavanine | 5.4 ±1.4 | 4.1 ±0.7 |
| At3g50210 | none | 20.9 ±1.7 | 0.78 ±0.15 |
| | Arginine | 21.3 ±3.7 | 1.75 ±0.43 |
| | Canavanine | 16.2 ±2.3 | 0.74 ±0.09 |

| | Arginine variable, 0.5 mM 2-oxoglutarate | | 2-Oxoglutarate variable, 10 mM homoarginine | | |
|---|---|---|---|---|---|
| | $A_{max}$ (nmol $s^{-1}$ $mg^{-1}$) | $K_M$ (mM) | $A_{max}$ (nmol $s^{-1}$ $mg^{-1}$) | $K_M$ (µM) | $K_I$ (mM) |
| Din11s | 3.1 ±1.4 | 6.0 ±5.1 | 5.2 ±0.9 | 30.1 ±9.4 | 17.6 ±9.6 |
| At3g49630 | n.d. | | 3.1 ±0.4 | 53.8 ±37.1 | 36.8 ±8.8 |
| At3g50210 | n.d. | | 19.0 ±7.2 | 69.7 ±23.2 | 35.3 ±20.1 |

All data are the average ±SD from 3 to 5 independent enzyme preparations

**Table S2: Arabidopsis T-DNA insertion lines used in this study**

| Line | NASC stock Nr. | Position of T-DNA | Reference |
|---|---|---|---|
| SALK_071632 | N571632 | 3rd intron of *Din11* | (Alonso et al., 2003) |
| GK-363E06 | N434806 | 1st exon of *Din11L*<br>5′-UTR of *Din11s* | (Kleinboelting et al., 2012) |
| SALK_118391 | N618391 | 4th intron of *At3g49630* | (Alonso et al., 2003) |
| SAIL_512_A07 | N821581 | 6th intron of *At3g49630* | (Sessions et al., 2002) |
| SALK_095112 | N595112 | 7th exon of *At3g49630* | (Alonso et al., 2003) |
| SALK_133126 | N633126 | 3rd exon of *At3g50210* | (Alonso et al., 2003) |
| SALK_098742 | N598742 | 1st intron of *At3g50210* | (Alonso et al., 2003) |

**Table S3: Sequences of primers used in this study**

| | |
|---|---|
| For cDNA insertion into pET24 (pET) or pENTR (pE) | |
| DIN11L-pET-f | gtactttcaaggtgctATGGTAATATATCATCGCAAAG |
| Din11s-pET-f | tgtactttcaaggtgctATGGTGACAGACTTCAAATCC |
| DIN11-pET-r | gtggtgctcgagtgcccataTTAACTGTTTTCCAC |
| DIN11L-pE-f | caggctttaaaggaacctATGGTAATATATCATCGCAAAG |
| DIN11s-pE-f | caggctttaaaggaacctATGGTGACAGACTTCAAATCC |
| DIN11-pE-r | gaaagctgggtctagataACTGTTTTCCACTAAATTTGC |
| 49630-pET-f | tgtactttcaaggtgctATGGCGACAAACTTCAAATCC |
| 49630-pET-r | ggtggtgctcgagtgctaacaTTAACTGTGTTCCACT |
| 49630-pE-f | caggctttaaaggaaccATGGCGACAAACTTCAAATCC |
| 49630-pE-NS-r | gaaagctgggtctagataACTGTGTTCCACTAAATTTTG |
| 50210-pET-f | tgtactttcaaggtgctATGGCGACGGACTTCAAGTCT |
| 50210-pET-r | ggtggtgctcgagtgccaTTACATAGCAAAGTTGGTC |
| 50210-pE-f | caggctttaaaggaaccatggcgacggacttcaag |
| 50210-pE-NS-r | gaaagctgggtctagatgCATAGCAAAGTTGGTCTGGA |
| Os_ODD33-pET-f | tgtactttcaaggtgctATGGGTTCTGACTTCAAGGC |
| Os_ODD33-pET-r | ggtggtgctcgagtgccaTTACATGACGAAGTTTGTCAAG |
| Gm_2-ODD-C23-pET-f | tgtactttcaaggtgctATGGCAACGGACTTTAGTTC |
| Gm_2-ODD-C23-pET-r | ggtggtgctcgagtgccaTTACAAATCAACAAAATTTGTGAGGACC |
| Ava_5009-pET-f | tgtactttcaaggtgcATGACAGTCTTACAACTTCCT |
| Ava_5009-pET-r | ggtggtgctcgagtgcctCTAAAGCACTTTTTGACG |
| ALDH7B4-pET-f | tgtactttcaaggtgctgctATGGGTTCGGCGAACAACGAG |
| ALDH7B4-pET-r | ggtggtgctcgagtgcCTAACCGAAGTTAATTCCTTGC |
| For Genotyping of T-DNA insertion lines | |
| Din11-f | TCAAGGGATTTGGATCACTCAC |
| Din11-r | TCGATCGAATGCATGCATCAC |
| At3g46930-f | CTGCTGGATACAGGTTTATTCCAC |
| At3g46930-r | CACGACAATACCCATTAGGCTC |
| At3g49630-f2 | GCTTGAGTGATGTTTCCAAGTAG |
| At3g49630-r2 | GAGGGATGGCAAAATCGTAG |
| At3g50210-f | CACCATGGCGACGGACTTCAAGTCT |
| At3g50210-r | AGGTTGGCCATTTGTGAACTCAG |
| SALK-LB | TTCGGAACCACCATCAAACAG |
| GABI-LB | AATAACGCTGCGGACATCTAC |
| SAIL-LB | TACCAATACATTACACTAGCATCTG |
| For generation and detection of genome editing sites | |
| gRNA-f | attgCAGATTGGTCATGGAATAT |
| gRNA-r | aaacATATTCCATGACCAATCTG |
| Din11-HRM-f | CTGATTATGGTTGGTGAGTGTG |
| Din11-HRM-r | CAACGATCGTTGATATAGCAGGT |
| At3g49630-HRM-f | TTGATTTTGGTCGGTGATTGTT |
| At3g49630-HRM-r | AGCCATCTAATCTAACTAATCGTGA |

Lower case letters indicate overlaps with vector sequences for cloning
